# Identification of Interaction Partners of Outer Inflammatory Protein A: Computational and Experimental Insights into How Helicobacter pylori Infects Host Cells

**DOI:** 10.1101/2024.03.03.583189

**Authors:** Sümeyye Akcelik-Deveci, Elif Kılıç, Nesteren Mansur Ozen, Emel Timucin, Yaren Buyukcolak, Sinem Oktem-Okullu

**Author notes:** Correspondence: Sinem Oktem-Okullu.

## Abstract

Adherence to the gastric epithelium is an essential feature of *Helicobacter pylori* for its colonization. Outer membrane proteins (OMPs) play a pivotal role in adherence potentiating the survival of the microbe in the gastric tissue. Among these proteins, Outer inflammatory protein A (OipA) is a critical protein that is known to help bacteria to colonize on the host gastric epithelial cell surface. Although the role of OipA in the *H. pylori* attachment and the association between OipA-positive *H. pylori* strains and clinical outcomes have been demonstrated, there is limited information on the structural mechanism of the OipA action in the adherence of *H. pylori* to the gastric epithelial cell surface. Our study utilizes experimental and computational methodologies to investigate the interaction partners of OipA on the gastric epithelial cell surface. Initially, we performed a proteomic analysis to decipher the OipA interactome in the human gastric epithelial cells using a pull-down assay of the recombinant OipA and the membrane proteins of the gastric epithelial cells. Proteomic analysis has revealed 704 unique proteins that interacted with OipA. We have further analyzed 16 partners of OipA using molecular modeling tools. Structural findings obtained from the prediction of the protein-protein complexes of OipA and candidate partners unraveled 3 human proteins whose OipA interactions could base an explanation about how *H. pylori* recruits OipA for adherence. Altogether, the findings presented here provide insights into novel mechanisms of *H. pylori* and host interactions through OipA, reflecting the potential of these mechanisms and interactions as therapeutic targets to combat *H. pylori* infection.

**Key points:**

- Outer membrane proteins (OMPs) are an emerging topic in bacterial infection.
- OipA is a candidate for an adherence-receptor network on the gastric epithelial cell surface with *H. pylori*.
- OipA interactome partners on gastric epithelial cell surfaces are valuable therapeutic targets for the *H. pylori* infection.

## Introduction

*Helicobacter pylori* (*H. pylori*) colonize the gastric epithelial cells of at least half of the world’s population and bacterial infection is associated with some gastric diseases, like chronic gastritis, and ulcer diseases. Persistent *H. pylori* infection with tissue damage can result in gastric cancer (Al-Maleki, Loke et al. 2017, Matsuo, Kido et al. 2017).

*H. pylori* infection is generally acquired during childhood and in the absence of treatment with antibiotics, it persists lifelong. Although the high prevalence of infections worldwide, most of the infected individuals remain asymptomatic for a long period. 15–20% of *H. pylori-infected* individuals develop at least one of the associated diseases at some point in their lives. The clinical outcome of *H. pylori* infection is believed to be influenced by multiple factors such as *H. pylori-related* virulence factors, host genetic predisposition, immune response, and environmental factors (Lima, de Lima et al. 2010) (Correa, Piazuelo et al. 2004) (Peek and Blaser 2002). It is essential to explain how different virulence factors contribute to the pathogenesis of *H. pylori* and its clinical consequences in order to develop a vaccine and create a more potent therapeutic strategy.

The outer membrane of *H. pylori* consists of two layers; the inner monolayer with only phospholipids and the outer monolayer with mainly outer membrane proteins (OMPs) which are resistant to the external environment (Qiao, Luo et al. 2014). OMPs have a variety of biological functions in *H. pylori* pathogenesis; maintaining the outer membrane structure, guaranteeing material transportation, and also playing an essential role in the process of contact with the host (Egan 2018) (Xu, Soyfoo et al. 2020).

Outer inflammatory protein A (OipA) is one of the most important outer membrane proteins (OMPs) that take a pivotal role in the virulence of *H. pylori*. OipA helps bacterial attachment to the gastric mucosa cells at the primary stage of the infection by establishing bacterial colonization. In the previous studies, it was shown that OipA-positive *H. pylori* strains have a stronger connection to the gastric mucosa than the strains that are OipA-negative. This highly antigenic protein increases the serum levels of interleukin-8 and the secretion of other inflammatory factors to cause neutrophil infiltration, aggravating the inflammation in the stomach. The binding of OipA to the host cell triggers an apoptotic cascade (Horridge, Begley et al. 2017) (Farzi, Yadegar et al. 2018) (Al-Maleki, Loke et al. 2017).

Some of the *H. pylori* OMP interaction partners on human gastric mucosa cell layers are known and their associated host cell responses have been gradually clarified (Matsuo, Kido et al. 2017). The role of OipA in the attachment of *H. pylori* to host cells has been confirmed. However, the interaction partners of OipA on gastric mucosa cell layers have not been defined and are the focus of this work.

In this study, we investigated the interaction partners of recombinant OipA in gastric cells by combining experimental and computational approaches. Initially, we cloned and heterologously expressed the OipA from *H. pylori* in *E. coli* cells. Purified recombinant OipA was then exposed to the o gastric cell membrane and surface proteins and the interactions were captured by immunoprecipitation. The resulting interactors were identified by mass spectroscopic methods and further filtered by a data-driven approach. Overall, sixteen proteins and their interaction with the extracellular region of OipA were delineated by extensive computational tools including protein structure prediction, rigid- and flexible protein-protein docking, and binding free energy calculations. Combined with experimental findings, computational findings showed both novel and partly identified OipA interactions as promising drug targets for combatting *H. pylori* infection of gastric cells.

## Materials and Methods

### *H. pylori* G27 Strain and Culture Conditions

*H. pylori* G27 strain was kindly donated by Prof. Dr. Anne Mueller from the University of Zurich, Institute of Molecular Cancer Research, Switzerland. The bacteria was grown on plates that consist of Colombia agar (OXOID), defibrinated horse blood, β-cyclodextrin (ChemCruz) solved with dimethyl sulfoxide, 200X (10 μg/mL Vancomycin (GeneMark), 5 μg/mL Cefsulodin (Koçak Pharma), 2.5 U/mL polymyxin B (ChemCruz)) and 1000X (5 μg/mL Trimetophrim (ChemCruz), 8 μg/mL Amphotericin B (Bristol-Myers Squibb), DMSO (SIGMA Life Science)) antibiotic-antifungal cocktails or 3-4 days at microaerophilic conditions (gas mixture composed of 5% oxygen, 10% carbon dioxide, and 85% nitrogen). The liquid cultures were grown on Brucella broth supplemented with 10% fetal bovine serum and 10 μg/mL vancomycin for 16-24 hours. For the storage of *H. pylori*, −80 ^°^C stock cultures freezing media was prepared that consists of brain heart infusion broth (BioShop^®^) containing 10% FBS (Gibco™) and 20% glycerol (SIGMA Life Science).

### DNA Extraction and Amplification of *oipA* Gene

Bacterial genomic DNA was extracted by using e Quick-DNA™ Miniprep Plus Kit (Zymo Research) according to the manufacturer’s instruction as the template. Briefly, bacterial cells harvested from the agar plate were resuspended in 1 ml of brucella broth and centrifuged to retain the pellet. The DNA concentration was measured with a NanoDrop 2000 (Thermo Fisher Scientific, United States). The *oipA* gene was amplified by polymerase chain reactions (PCRs) with forward primer OipA F 5’-CATTAAGCGGTGGTTTTGTG-3’ and reverse primers OipA R 5’-AGCCAACTAAAGAGCGGTAA-3’ The PCR reaction was performed in a total volume of 25 μl containing Pfu DNA polymerase (GeneMark), forward and reverse primers (0.75μl each), 2 μl DNA template, 2.5 μl dNTP and 16.5 μl nuclease-free water. The amplification was as follows: denaturation at 95^°^C for 5 min, 39 cycles of denaturation at 95^°^C for 30 s, annealing at 59.2^°^C for 1 min, extension at 72^°^C for 2 min, and a final extension at 72^°^C for 7 min. PCR products were analyzed under UV light with ChemiDoc (Thermo Fisher Scientific, United States) after electrophoresis in a 1% agarose gel.

### Gene Cloning and Recombinant Protein Expression

For *oipA* gene cloning, extracted *H. pylori G27* DNA samples were subjected to PCR with C-His oipA forward primer 5’-AGAAGGAGATATAACTATGATGAAAAAAGCTCTCTTACT-3’ and reverse primer 5’-GGAGATGGGAAGTCATTAATGATGGTGATGGTGGTGATGTTTGTTTTTAAAGTT-3’ that are designed to add His6 peptide sequence to the *oipA* gene. The amplified PCR product of the C-His *oipA* gene fragment was cloned into vector pLATE11 using aLICator LIC Cloning and Expression Kit 1 (Thermo Fisher Scientific, United States) by following the kit’s instructions. The transfection of constructed vector (pLATE11+C-His oipA) was transfected into the competent *E. coli BL21 (DE3)* strain with the heat shock transfection method. After the appearance of transfected colonies on a selective agar plate, the transfection was validated by colony PCR and the chosen colony was used to produce of C-His oipA protein.

### Purification of Recombinant C-His-OipA Protein

Isolated total protein from *E. coli BL21 (DE3)* strain was loaded onto the SDS PAGE gel and target protein band (between 25-35 kda) was eluted from the gel by passive elution method (Kurien and Scofield 2012). C-His-OipA protein was purified by using Dynabeads^™^ His-Tag Isolation and Pulldown magnetic bead system (Thermo Fisher Scientific, United States). His-tagged OipA protein isolation procedure was carried out according to kit instructions. Briefly, protein solution was prepared with the 1X binding and wash buffer that includes 50mM sodium phosphate, pH 8.0, 300 mM NaCl, and 0.01% Tween^™^-20. Then the prepared mixture was incubated with cobalt magnetic beads at 4 ^°^C for 10 minutes. After incubation unbound proteins were washed away and bounded proteins were eluted by using a His elution buffer solution containing 300 mM Imidazole, 50 mM Sodium phosphate, pH 8.0, 300 mM NaCl, and 0.01% Tween^™^-20.

For the refolding of purified recombinant C-His OipA protein dialysis method was used in which chemically denatured protein is refolded to sufficiently decrease the denaturant concentration and allow protein refolding. The volume of the refolding buffer that includes 0.1 mM DTT and 20mM Tris HCl pH 8.5 was used up to 200 times the purified recombinant protein volume. Recombinant purified protein was transferred to the dialysis membrane for which the length was calculated following dialysis supplier company instructions. The solution was changed 2 times with an interval of 3 hours. The solution was then replaced with a cold solution containing only 20mM Tris-HCl twice at 3-hour intervals at 4ºC, and the protein in the dialysis membrane was centrifuged at 10,000g for 10 minutes at 4ºC.

### Isolation of Membrane Proteins

The AGS cell line derived from the human gastric adenocarcinoma (ATCC® CRL-1739™) was routinely cultivated at 37 ^°^C and 5 % CO_2_ in the RPMI 1640 medium (Gibco). The growth media contained 10% fetal bovine serum (Gibco) and 1% penicillin/streptomycin (Gibco). The membrane proteins of AGS cells were isolated with the Mem-PER™ Plus Membrane Protein Extraction Kit (Thermo Fisher Scientific, United States). Briefly, 5×10^6^ AGS cells were lysed with the kit’s detergent, and during the centrifugation steps, the cell’s hydrophobic end hydrophilic protein phases were distributed. To maintain the 3D structure of protein mild detergents were removed by using Pierce™ Detergent Removal Spin Column, 0.5 mL kit (Thermo Fisher Scientific, United States). After taking the hydrophobic fraction of the AGS cell proteins the protein concentrations were determined by Bicinchoninic acid assay.

### Histidine-Pull-down Interaction Analysis

Dynabeads™ His-Tag Isolation and Pulldown magnetic bead solution (Thermo Fisher Scientific, United States) kit was used for the detection of the binding partner of OipA. The refolded Histidine-tagged OipA protein was prepared with the 1 X binding and wash buffer and then incubated with cobalt-based magnetic beads as described in the purification step. After this step unbound proteins were washed away and then isolated AGS membrane proteins in the pull-down buffer (3.25 mM Sodium phosphate, pH 7.4, 70 mM NaCl, 0.01% Tween™-20) were mixed with bounded OipA proteins on magnetic beads and incubated for 1 h at 4 ^°^C. After incubation the protein mixture was eluted by His elution buffer. The eluted solutions were analyzed by LC-MS/MS.

### LC-MS/MS Analysis

“In solution digestion’’ was performed as described previously (Polat, Karayel et al. 2015). Briefly, cell pellets were lysed in a buffer containing 8M urea, 50 mM ammonium bicarbonate and protease inhibitor tablets. Following ultrasonication, lysates were centrifuged in 14000g at 4^°^C for 45 minutes and supernatants were collected. Protein samples were reduced with DTT and alkylated using iodoacetamide. Prior to trypsin addition, samples were diluted 8-fold using 50 mM ammonium bicarbonate and incubated with trypsin overnight. Next morning, reaction was stopped by acidification with 10% formic acid and digested peptides were desalted using solid phase extraction (SPE).

Digested peptides were analyzed with 90-minute gradients in C18 nanoflow reversed-phase HPLC (Dionex Ultimate 3000,3500 RSLC nano, Thermo Scientific) combined with orbitrap mass spectrometer (Q Exactive Orbitrap HF, Thermo Scientific). Scan parameters for MS1 were; 70 000-resolution; AGC 3e6; Max IT 60 ms; Scan range 400-1500 m/z and for MS2; 17 500-resolution; AGC:5e4; Max IT:60 ms; Top 15; Isolation window:2.0 m/z; NCE: 26. Raw files were processes in Thermo Scientific Proteome Discoverer 2.3 software using Sequest search engine and human Uniprot database (Release 2016).”

### Protein-protein docking

The full-length of OipA structure was predicted by Alphafold using the open-source Colab notebook (Jumper, Evans et al. 2021). Predicted structure was locally assessed based on pLDDT scores and predicted aligned error. The extracellular domain of the OipA (OipA ED) was analyzed in rigid-body docking by Cluspro algorithm. Overall a total of 20 proteins that were enlisted after MS analysis were analyzed in this docking round. Ten of the top-scoring rigid complexes were superimposed to identify the dominant binding interface of the rigid docking runs. The amino acids that contoured at least 50% of the top-scoring complexes were used as an input to the flexible docking method (van Zundert, Rodrigues et al. 2016). HADDOCK runs were performed for each complex by introducing flexibility to the interconnecting loops at the identified interface. The complexes were refined after a final iteration in water. The first representative of the top HADDOCK cluster was selected for each run. The final complexes were selected based on docking scores.

## Results

### Identification of OipA Interacting Partner on Human Gastric Epithelial Cell Surface

According to the optimization studies C-His *oipA* cloned colonies were observed in 1: 7 vector: insert molar ratio and T4 DNA polymerase activity at 50^°^C for 30 seconds. After the transformation, depending on the Sanger sequencing confirmation the recombinant pLATE11-C-His oipA plasmid was transferred to *E. coli* BL21 (DE3) competent cell. 1 mM IPTG at 37^°^C for 4 hours was used for the induction of protein as a result of serial optimization studies. Results of purification of recombinant C-His-OipA protein by magnetic bead were confirmed by western blot analysis. To identify the OipA interacting partner, the pull-down assay with 80 μl nickel magnetic beads of bait and prey protein at a ratio of 1:4. 160 μl of C-His OipA at 10 ng/μl concentration and 640 μl AGS membrane protein were used (Figure 1 (a-b)). AGS membrane proteins were mixed with immobilized refolded C-His OipA protein into cobalt magnetic beads. The obtained protein samples were analyzed with LC-MS/MS (Figure 1c).

**Figure 1.**
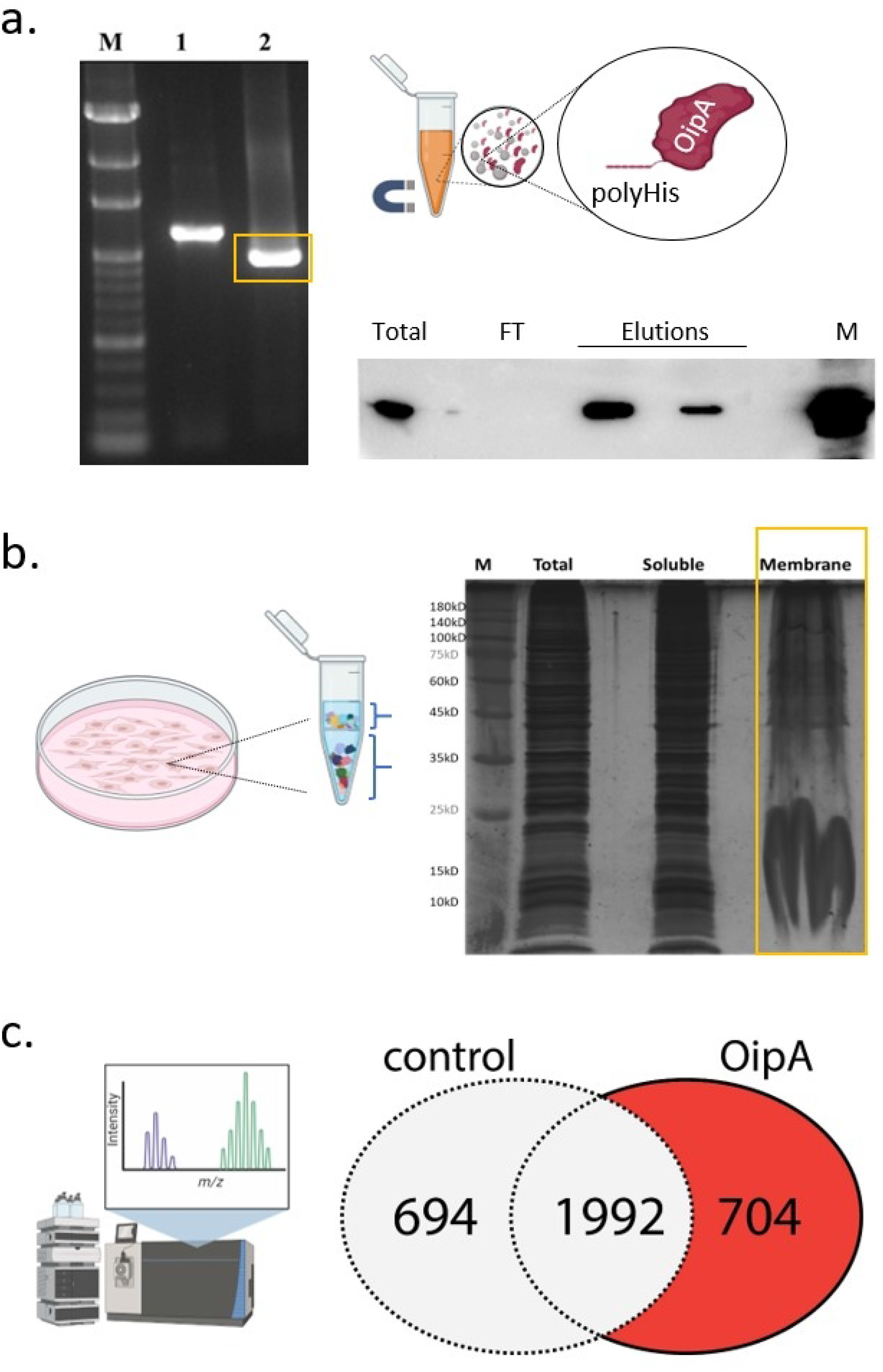
(a) Molecular cloning of the OipA gene (left) from *H. pylori* g27 genome into the aLICator LIC Cloning and Expression system. M: 100bp marker (GeneMark), 1: PCR product by priming external regions of *oipA* gene, 2: Full-length *oipA* gene (964bp). After expression of the *oipA* plasmid in the *E. coli* BL21 (DE3), recombinant OipA was purified by cobalt-coated magnetic beads. Western blot analysis of the purified OipA using anti-His antibody (CellSignalling) (b) Results of membrane protein extraction from AGS cells (Mem-PER^TM^-Thermo) were shown on silver-stained SDS-PAGE gel. (c) Pull-down analysis of OipA interacting proteins from the membrane lane in (b) by LC-MS/MS. Venn diagram shows the number of identified interactors in OipA overexpressing samples and control.

### Identification of OipA Interacting Proteins

To identify OipA interacting proteins, we have selected the proteins that appeared in all three OipA over-expressing samples but in the control samples. A total of 704 different proteins were listed and analyzed using subcellular compartments. The proteins that were localized to the cell membrane either by anchoring to the outer membrane or by spanning the bilayer were reported as OipA interacting proteins (Table S1). Proteins localized to other compartments were not selected for structural modeling; however, elucidation of the atomistic details of their interactions with OipA would also be interesting given the observation that they were listed as OipA interacting proteins in all three replicates.

### Prediction of the full-length OipA Structure

The structure of OipA or any other close relatives has not been experimentally characterized yet because they reside in the outer membrane of *H. pylori*. However, AlphaFold2 (AF2), which predicts highly accurate protein structures, could be implemented for prediction of the OipA structure. This prediction could further equip us with the complex structural models formed between OipA and its partners.

Due to the absence of another structure with an apparent homology to OipA, comparative modeling techniques would likely fail or produce unreliable predictions for the OipA structure. As such, we have tried both homology modeling and threading approaches to predict the 3D structure of *H. pylori*’s OipA and yet the results were not satisfactory to meet the structure-function relationship (Figure 2). Given the advance of the robust deep learning method, AlphaFold (AF) (Tunyasuvunakool, Adler et al. 2021) (Jumper, Evans et al. 2021) for the prediction of protein structures without any structural templates, we have employed AF for the prediction of the full-length OipA structure. Results of AF prediction were given in Figure 2. At this point, we are well aware that, AF or not, computational models reflecting predictions would be problematic as would thus blindly relying on such models would also be problematic. However, we report the following points for this particular prediction of OipA structure which agrees that the model in fact could represent the physical structure. First based on the pLDDT scores which reflect the confidence of the AF prediction, the first-ever OipA structure was largely predicted with a high accuracy i.e. high pLDDT score. Second the resulting 3D fold is a beta-barrel with 5 beta strands. Barrel shape when fitted in between a membrane bilayer we observed that the barrel was completely embedded in the bilayer. Further, the barrel was lined up with highly hydrophobic amino acids which is another indication of the reliability of the prediction. Thus, the resulting OipA structure could be further examined to address whether or not the listed interacting partners from the mass spectroscopy analysis are of the form a stable complex.

**Figure 2:**
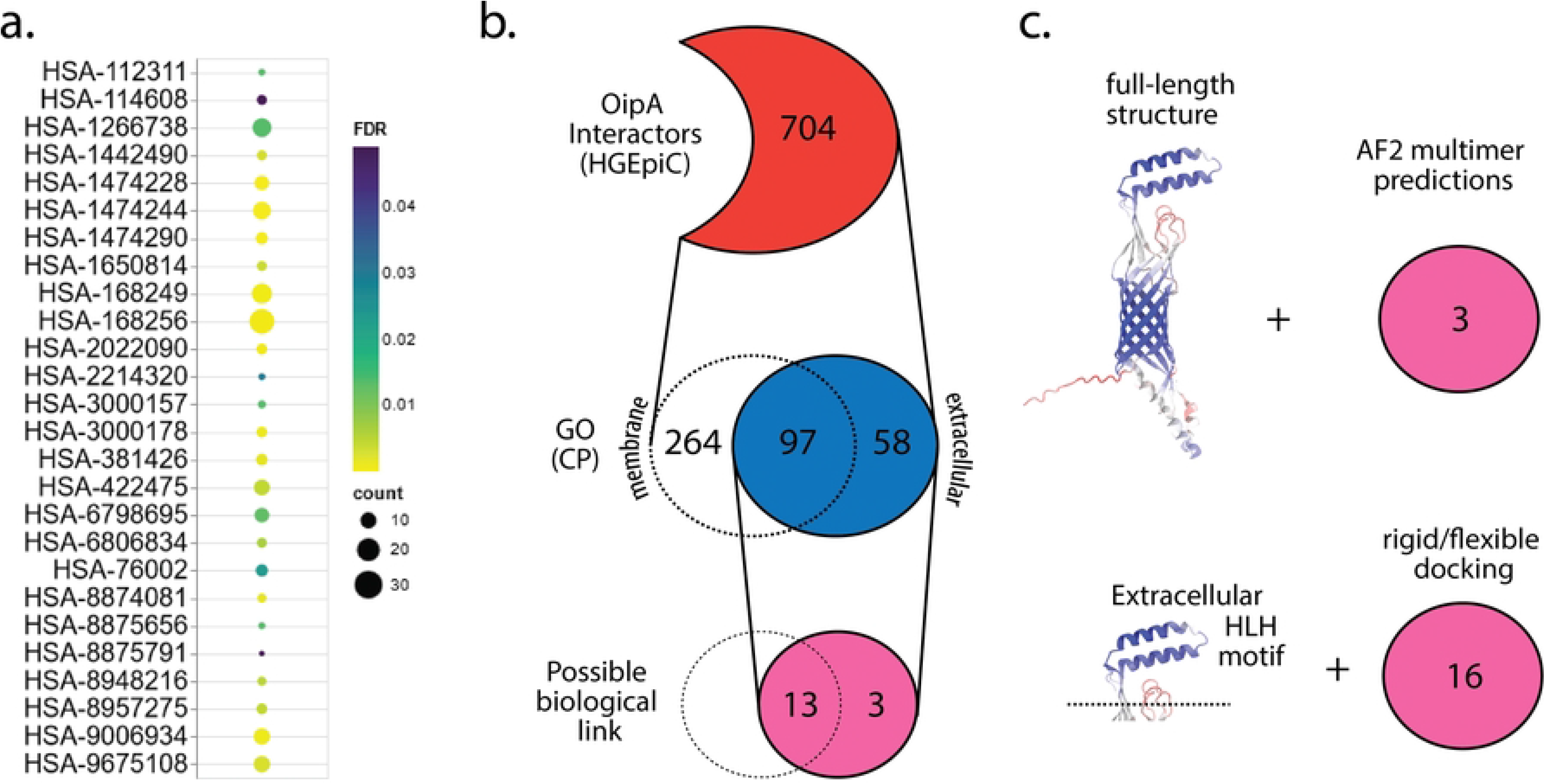
(a) shows the Reactome pathways for the proteins that are found interacting with OipA and have expected cellular compartment GO annotation. Color-scale shows the corrected p-values for the analysis. Table S1 enlists the enriched pathways shown here. (b) OipA interactors were screened based on their GO cellular compartment annotation and the proteins that are either secreted or have extracellular domains/regions were considered. Biological link was established if there has been any previous study linking the identified partner and OipA and/or *H. pylori*. (c) Shows the structural analysis conducted in this study. After full-length structure prediction of OipA, either the full-length structure or the small extracellular portion was used in prediction of the complex interactions that were identified in proteomics analysis.

Aiming to understand the role of OipA in host-cell interactions, we have limited our focus to the extracellular proteins or proteins having an extracellular domain. Further, we also focused on the alpha-turn-alpha motif of the OipA structure which extends to the extracellular region. We referred to this OipA as the extracellular domain of OipA (ECD). Among 704 number of proteins ensuring the criteria provided above, we have reduced our list to 16 through a thorough literature search which confirmed that the selected proteins were previously implicated in host interactions of other microbes or *H. pylori*. Three-dimensional structure of OipA complexes with the selected 16 proteins predicted by two protein-protein docking algorithms. The resulting OipA complexes were visualized (Figure 4). Rigid-body predictions of these complexes formed showed that the bound complex consistently shared the same interface, suggesting that a convergence of rigid-body results. Next, the flexible docking runs were carried out to refine the docked complexes shown in Figure 4. Docking scores obtained from the second round of flexible docking was listed in Table 1.

**Table 1.** List of possible OipA interactors that were analyzed in molecular docking.

| UniProt | Gene name | Structure (source) | Score | Ref* |
| --- | --- | --- | --- | --- |
| P00918 | CA-II | 1BCD (PDB) | -133.4±1.5 | (Campestre, De Luca et al. 2021),(Bury-Moné, Mendz et al. 2008) |
| O14514 | AGRB1 | ADGRB1-F1 (AF) | -82.7±1.7 | (Billings, Lee et al. 2016) |
| Q16853 | VAP-1 | 4BTX (PDB) | -131.9±0.3 | (Svensson, Hansson et al. 2009) |
| P05452 | TETN | 1HTN (PDB) | -69.1±1.3 | (Pantzar, Ljungh et al. 1998) |
| Q15828 | CYTM | 4N6L (PDB) | -65.8±2.7 | (Szulc-Dąbrowska, Bossowska-Nowicka et al. 2020) |
| Q9UQC9 | CLCA2 | AF-Q9UQC9-F1 (AF) | -119.0±6.8 | (Liu and Shi 2019),(Loewen and Forsyth 2005) |
| P69905 | HBA1 | 2DN1 (PDB) | -121.4±2.4 | (Bowden, Chan et al. 2018) |
| P20338 | RAB4A | 2BMD (PDB) | -93.6±2.9 | (Zhang, Fan et al. 2005) |
| O14745 | NHRF1 | 4Q3H (PDB) | -90.4±1.7 | (Martin, Barbieri et al. 2020) |
| Q12805 | FBLN3 | Q12805-F1 (AF) | -93.1±0.6 | (Min, Zhao et al. 2017) |
| P08581 | MET | 2UZX (PDB) | -132.1±2.3 | (Ito, Tsujimoto et al. 2020), (Niemann, Jäger et al. 2007) |
| Q9NVZ3 | NECP2 | Q9NVZ3-F1 (AF) | -90.8±2.2 | (Ritter, Philie et al. 2003) |
| P14210 | HGF | 4O3T (PDB) | -103.4±0.8 | (Ito, Tsujimoto et al. 2020), (Xie, Yang et al. 2017),(Franke, Müller et al. 2008) |
| O00560 | SDCB1 | 1NTE (PDB) | -68.6±0.4 | (Hayashida, Amano et al. 2011) |
| P55290 | CAD13 | 2V37 (PDB) | -64.9±4.0 | (Connell, Meade et al. 2013) |
| P27361 | MAPK 3 | 4QTB (PDB) | -118.6±5.1 | (Meyer-ter-Vehn, Covacci et al. 2000), (Ellington, Elhofy et al. 2001) |
| *HADDOCK score is in kcal/mol. |  |  |  |  |

We also focused on the other interactions of these partners. Strikingly, we spotted MET protein which was previously shown to interact with *L. monocytogenes* internalin protein for attachment. Given its previous involvement in other host-microbe interactions, MET could also interact with the extracellular domain of OipA and the refined complex model shown in Fig. 5a. is the first prediction of such interaction of OipA and thus has paramount importance for therapeutic strategies against *H. pylori* infections.

We also analyzed the interaction between the full-length OipA and its three partners namely, HGF, MET and AGRB1. For modeling full-length complex structure, we have utilized AF2 multimer (Mirdita, Schutze et al. 2022, Zhu, Shenoy et al. 2023). Resulting complexes were somewhat different from the predictions made for the extracellular helix-loop-helix motif of the OipA (Figure 5a-MET, 5b-HGF, 5c-AGRB1). Particularly the binding of full-length OIPA to MET took place from the extracellular domain of MET (Figure 5a). On the hand, both predictions for the HGF complex overlapped with each other marking a small cleft on the HGF structure (Figure 5b). However, the predictions did not show the same OipA surface as such for the full length prediction, the binding interface consisted of the small extracellular helix-loop-helix motif along with the outer mouth of the β-barrel structure (Figure 5b). Overall, the full-length predictions in comparison with the docking of the small helix-loop-helix motif of OipA provided insights into the possible complexes of OipA with several key proteins that could base a structural explanation to OipA’s role in adhesion.

## Discussion

Despite gastric peristalsis, *H. pylori* demonstrate powerful interaction with gastric epithelial cell (Posselt, Backert et al. 2013) and has developed some mechanisms to cause infection in the harsh acidic stomach environment. The pathogenesis of *H. pylori* infection and disease outcome are explained by the relationship between host genetics, environmental and bacterial virulence factors (Braga, Batista et al. 2019). The virulence factors take an important role for adherence, colonization, and activation of the host immune response. Especially, adherence of *H. pylori* to the mucus layer of the gastric epithelium has an significant role in the initiation of colonization that continues with the infection (Oleastro and Ménard 2013). The extraordinary large outer membrane protein (OMP) family of the bacterium includes proteins specifically involved in attachment (Kalali, Mejías-Luque et al. 2014). The interaction of bacterial outer membrane proteins with cellular receptors protects the bacteria from mechanisms like acidic pH, mucus, and exfoliation of stomach (Kalali, Mejías-Luque et al. 2014, Kao, Sheu et al. 2016). OipA is one of the important outer membrane protein that attach to gastric epithelial cells in bacterial pathogenesis. The functional status of OipA are determined by a slipped strand mispairing mechanism. The patients infected with *H. pylori* has a functional OipA have a higher risk of developing gastric cancer. OipA is an important virulence factor as it provides a stronger effect by interfering with the signal pathways activated by CagA/T4SS. OipA mediates a tight attachment between bacteria and gastric epithelial cells through this strong effect that occurs either indirectly or by binding to the surface protein, in colonization (Posselt, Backert et al. 2013). The functions and regulated signals of OipA in epithelial cells have been explained, but the surface protein with which OipA binding interacts in epithelial cells has not been identified. Thus, we performed this study to find the interaction partner of OipA, and identified proteins from AGS cell lines by pull down assay followed by LC-MS/MS analysis.

Proteomic analysis showed 704 unique binding partners of OipA in gastric epithelial cells. Inspection of this partner list led to a collection of a small set of OipA binders that were selected based on subcellular localization and biological relevancy. This small set includes Hepatocyte growth factor (HGF), Hepatocyte growth factor receptor (MET), Carbonic anhydrase II (CA-II), Adhesion G protein-coupled receptor B1 (AGRB1), Vascular adhesion protein 1 (VAP-1), Tetranectin, (TETN), Cystatin-M (CYTM), Calcium-activated chloride channel regulator 2 (CLCA2), Hemoglobin subunit alpha (HBA1), Ras-related protein Rab-4A (RAB4A), Na(+)/H(+) exchange regulatory cofactor (NHRF1), EGF-containing fibulin-like extracellular matrix protein 1 (FBLN3), Adaptin ear-binding coat-associated protein 2 (NECP2), Syntenin-1 (SDCB1), Cadherin-13 (CAD13), Mitogen-activated protein kinase 3 (MAPK3). All 16 protein partners were extensively analyzed by molecular modeling methods, providing 16 distinct atom-resolution complex structures of OipA.

The quality of the computed structures that were not derived from any experimental data should be properly evaluated. Among the recent artificial intelligence methods that were developed to predict protein structure, most of the tools provide a mean of quality evaluation. The method that was employed here, AF2, essentially uses a metric for assessment of the reliability of the predicted Cα positions (Mariani, Biasini et al. 2013), which is called the predicted local difference distance test (pLDDT) score. Given the availability of such a measure at the residue-level, one could assess the quality of the predicted OipA backbone structure. From this regard, we considered the pLDDT scores of each amino acid for the OipA structure and found out that the full-length AF2 structure of OipA having most of the amino acids with the pLDDT scores higher than 70 (Figure 3). Especially, the pLDDT based scores were much higher at the core β-barrel region while predictably the confidence scores of the loop regions were lower than the threshold of 50. Nonetheless, we underscore that the first structural modeling of the OipA structure would open new doors to understand the molecular mechanism of the protein in the pathogenesis of *H. pylori*.

**Figure 3.**
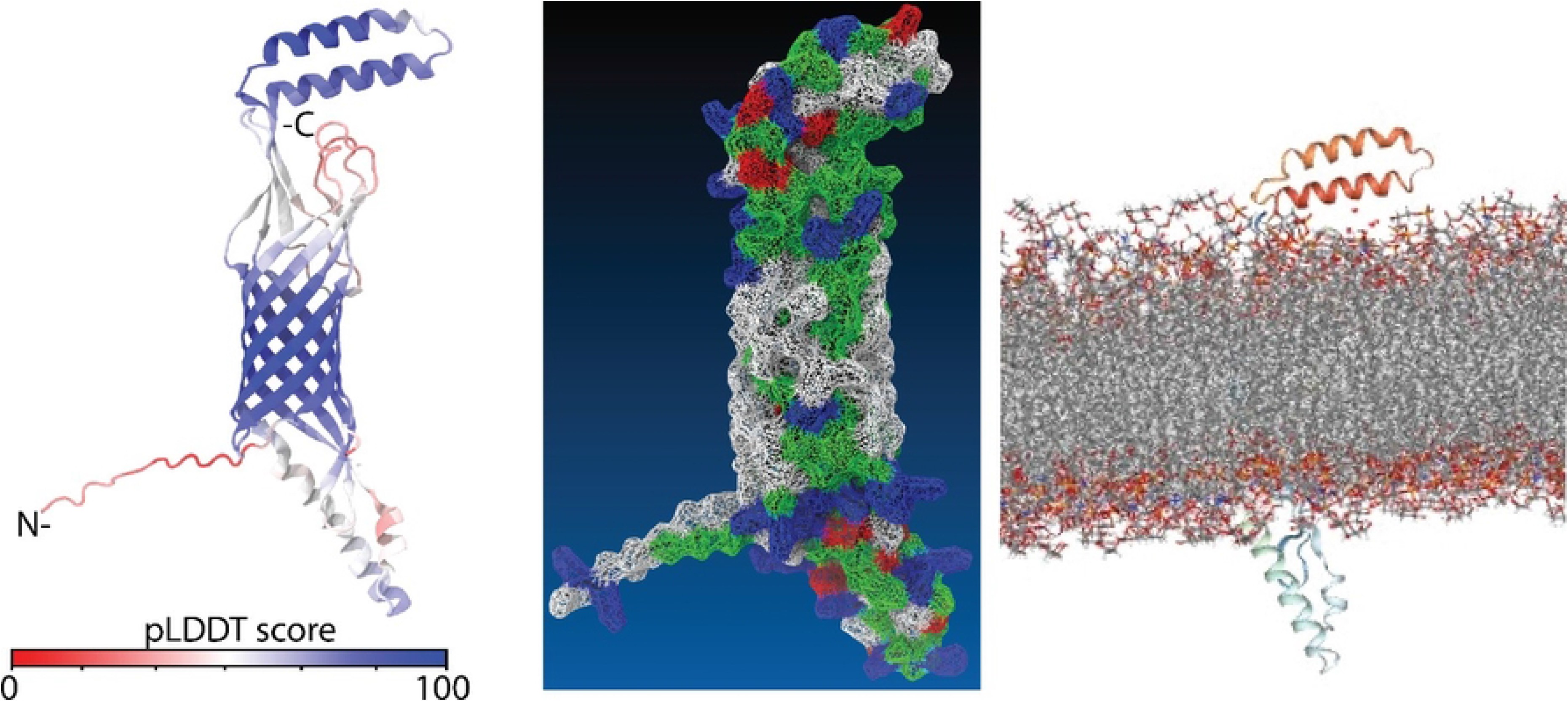
AF prediction of OipA structure: left shows the backbone conformation colored according to the pLDDT score of the prediction, the middle panel shows the surface contour of the same structure (white: hydrophobic, blue: basic, red: acidic, green: polar amino acids), right shows the structure embedded in a membrane bilayer formed POPE lipids. Membrane embedding was performed by the CHARMM-GUI web server.

Structure-based modeling also unraveled the novel complexes of OipA and Hepatocyte growth factor (HGF) as well as Hepatocyte growth factor receptor (MET) that were tightly linked with infectious diseases and chronic inflammation. Essentially, HGF drives mitogenesis, motogenesis, and morphogenesis in a variety of tissues, including epithelial cells upon binding to the receptor tyrosine kinase c-Met. The HGF-Met axis has been linked to several infectious diseases as such several pathogens have been found to hijack the HGF-Met system by down regulating Met signaling altering the expression levels of HGF or Met through the use of their pathogenic factors. Hence, real-time detection of Met activation and monitoring of HGF and Met expression is useful for detection of viral infections. From this perspective, the HGF-Met system stood out as therapeutic targets to fight infectious disorders (Imamura and Matsumoto 2017). Preclinical research on injury/disease models, such as acute tissue injury, chronic fibrosis, cardiovascular illness, and neurodegenerative disorders, have been carried out to address the therapeutic importance of HGF. HGF/c-Met signaling pathway was also associated with the polarity of the epithelial cell that affects the cell susceptibility to pathogenic microorganism (Wu, Billings et al. 2001). Importantly, HGF/c-Met signaling pathway is targeted by one of the important virulence factors, CagA resulting in enhancement of cell scattering which is an indicator of the motogenic response of *H. pylori*-infected gastric epithelial cells (Furge, Zhang et al. 2000, Churin, Al-Ghoul et al. 2003, Franke, Müller et al. 2008). Furthermore, another Gram negative human pathogen *L. monocytogenes* also uses HGF-Met axis for invasion of host cells. Particularly for this pathogen, a structural study has revealed that a crystal complex structure between internalin, a InlB protein of the pathogen and MET (Braun, Dramsi et al. 1997). Comparison of this structure with our OipA-MET complexes showed a partial overlap in the binding interface of both of the models (Figure 5a). Altogether, the importance of the pivotal role played by the HGF-Met signaling pathway in the progression of infectious diseases and our proteomic analysis suggested a binding activity of the recombinant OipA towards the MET on the gastric epithelial cell surface. Our structural predictions have further implied that this interaction occurs from a similar MET surface that were utilized by *L. monocytogenes*. Thus, we consider that along with the established literature on the HGF/Met and CagA interaction, our findings reported novel insights into how *H. pylori* infects its host though the HGF/Met-OipA interaction consolidating the potential of HGF/Met as a therapeutic target for this infection.

**Figure 4:**
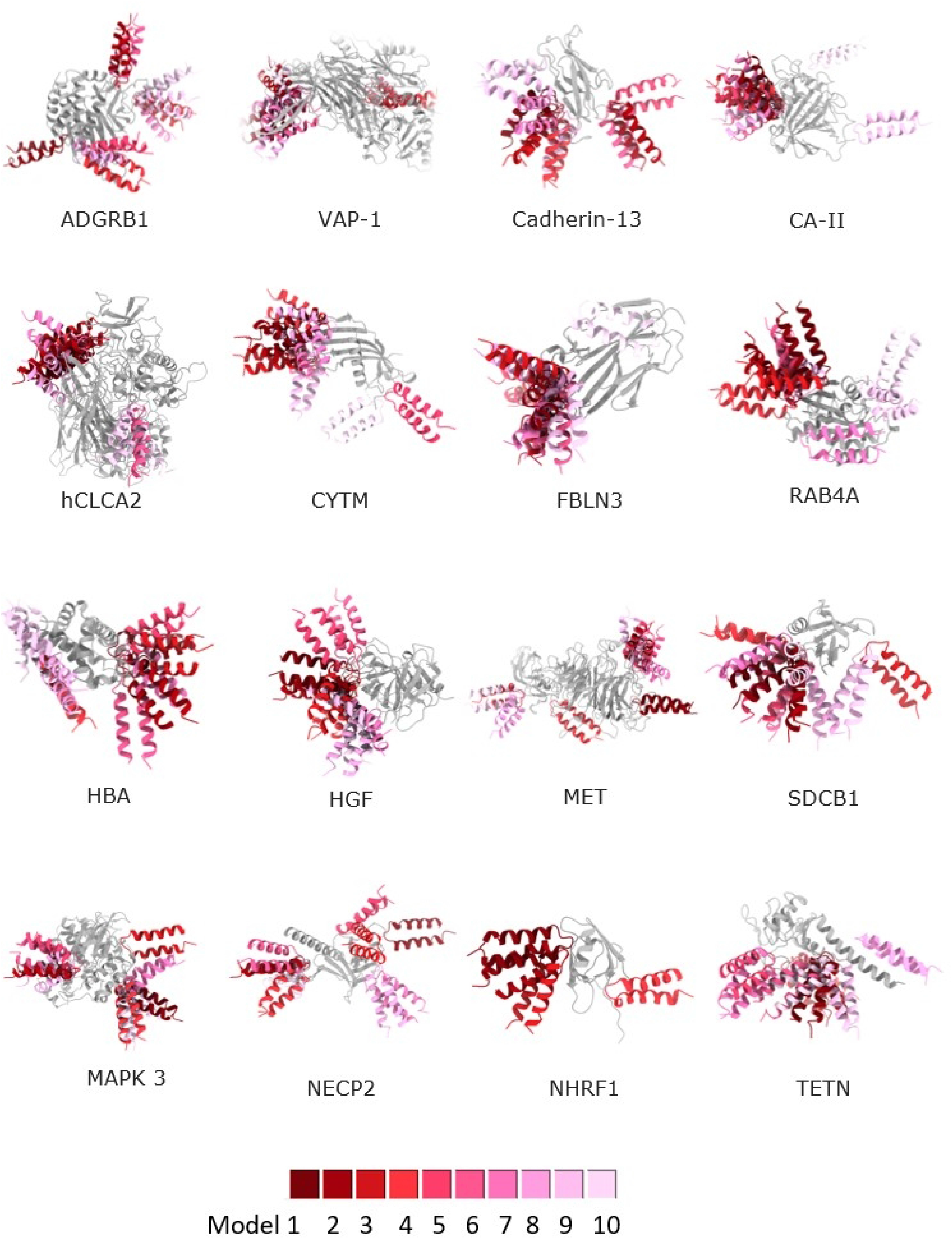
First docking analysis of the Oipa interacting partners listed in Table 1 and the extracellular domain of Oipa. Analysis is done by ClusPro and top ten high scoring models were illustrated.

**Figure 5:**
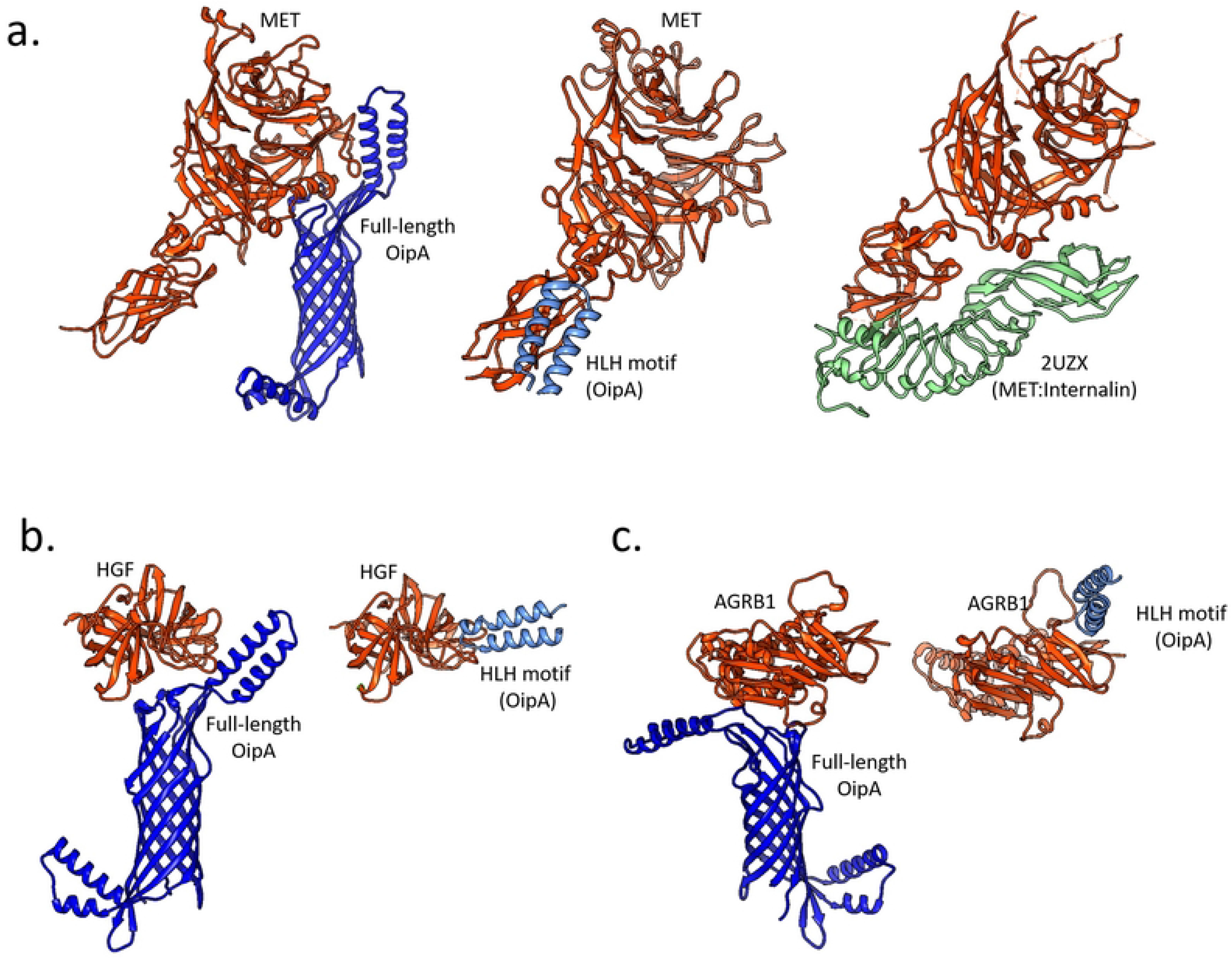
Results of OipA complexes that were predicted either using the full-length structure (AF2) or the helix-loop-helix motif (docking). (a) Panel shows the predicted complexes with the MET. The crystal structure that captured MET and L. monocytoge*nes* internalin protein. (b) Panel shows the complexes formed with HGF and (c) shows formed with AGRB1.

Another binding partner of OipA, Adhesion G protein-coupled receptor B1 (AGRB1) which is a phosphatidylserine receptor was identified to mediate binding and engulfment of particularly Gram-negative pathogens (Billings, Lee et al. 2016) (Das, Owen et al. 2011). Here, we also identified how this protein could interact with OipA, using both full-length and partial OipA structures. These analyses implied that the AGRB1 interaction was likely to take place on the outer part of OipA as the full-length model showed that the AGRB1 attached to the extracellular part of the OipA, in fact showing a flexed helix-loop-helix motif (Figure 5c).

The binding of OipA protein and candidate interaction partner is an important step in the attachment of *H. pylori* to the gastric mucus layer. The interaction partners can be targeted to prevent this infection. Hence, we conclude that the reported interaction partners of OipA that were determined by a proteomic analysis could be highly promising for future studies aiming to examine the molecular mechanism(s) of OipA in *H. pylori* infection.

## Funding

This work was supported by grants from the The Scientific and Technological Research Council of Turkey with the project number 217S384.

## Data availability

All data will be made available from the corresponding author upon request.

## Declarations

### Ethical approval

This article does not contain any studies with human participants or animals performed by any of the authors.

### Conflict of interest

The authors declare no competing interests.

## Notes

### Competing Interest Statement

The authors have declared no competing interest.

## References

Al-Maleki, A. R., M. F. Loke, S. Y. Lui, N. S. K. Ramli, Y. Khosravi, C. G. Ng, G. Venkatraman, K. L. Goh, B. Ho and J. Vadivelu (2017). “Helicobacter pylori outer inflammatory protein A (OipA) suppresses apoptosis of AGS gastric cells in vitro.” Cell Microbiol 19(12).

Billings, E. A., C. S. Lee, K. A. Owen, R. S. D’Souza, K. S. Ravichandran and J. E. Casanova (2016). “The adhesion GPCR BAI1 mediates macrophage ROS production and microbicidal activity against Gram-negative bacteria.” Sci Signal 9(413): ra14.

Bowden, C. F. M., A. C. K. Chan, E. J. W. Li, A. L. Arrieta, L. D. Eltis and M. E. P. Murphy (2018). “Structure-function analyses reveal key features in Staphylococcus aureus IsdB-associated unfolding of the heme-binding pocket of human hemoglobin.” J Biol Chem 293(1): 177–190.

Braga, L., M. H. R. Batista, O. G. R. de Azevedo, K. C. da Silva Costa, A. D. Gomes, G. A. Rocha and D. M. M. Queiroz (2019). “oipA “on” status of Helicobacter pylori is associated with gastric cancer in North-Eastern Brazil.” BMC Cancer 19(1): 48.

Braun, L., S. Dramsi, P. Dehoux, H. Bierne, G. Lindahl and P. Cossart (1997). “InlB: an invasion protein of Listeria monocytogenes with a novel type of surface association.” Mol Microbiol 25(2): 285–294.

Bury-Moné, S., G. L. Mendz, G. E. Ball, M. Thibonnier, K. Stingl, C. Ecobichon, P. Avé, M. Huerre, A. Labigne, J. M. Thiberge and H. De Reuse (2008). “Roles of alpha and beta carbonic anhydrases of Helicobacter pylori in the urease-dependent response to acidity and in colonization of the murine gastric mucosa.” Infect Immun 76(2): 497–509.

Campestre, C., V. De Luca, S. Carradori, R. Grande, V. Carginale, A. Scaloni, C. T. Supuran and C. Capasso (2021). “Carbonic Anhydrases: New Perspectives on Protein Functional Role and Inhibition in Helicobacter pylori.” Frontiers in Microbiology 12.

Churin, Y., L. Al-Ghoul, O. Kepp, T. F. Meyer, W. Birchmeier and M. Naumann (2003). “Helicobacter pylori CagA protein targets the c-Met receptor and enhances the motogenic response.” J Cell Biol 161(2): 249–255.

Connell, S., K. G. Meade, B. Allan, A. T. Lloyd, T. Downing, C. O’Farrelly and D. G. Bradley (2013). “Genome-wide association analysis of avian resistance to Campylobacter jejuni colonization identifies risk locus spanning the CDH13 gene.” G3 (Bethesda) 3(5): 881–890.

Correa, P., M. B. Piazuelo and M. C. Camargo (2004). “The future of gastric cancer prevention.” Gastric Cancer 7(1): 9–16.

Das, S., K. A. Owen, K. T. Ly, D. Park, S. G. Black, J. M. Wilson, C. D. Sifri, K. S. Ravichandran, P. B. Ernst and J. E. Casanova (2011). “Brain angiogenesis inhibitor 1 (BAI1) is a pattern recognition receptor that mediates macrophage binding and engulfment of Gram-negative bacteria.” Proc Natl Acad Sci U S A 108(5): 2136–2141.

Egan, A. J. F. (2018). “Bacterial outer membrane constriction.” Mol Microbiol 107(6): 676–687.

Ellington, J. K., A. Elhofy, K. L. Bost and M. C. Hudson (2001). “Involvement of mitogen-activated protein kinase pathways in Staphylococcus aureus invasion of normal osteoblasts.” Infect Immun 69(9): 5235–5242.

Farzi, N., A. Yadegar, H. A. Aghdaei, Y. Yamaoka and M. R. Zali (2018). “Genetic diversity and functional analysis of oipA gene in association with other virulence factors among Helicobacter pylori isolates from Iranian patients with different gastric diseases.” Infect Genet Evol 60: 26–34.

Franke, R., M. Müller, N. Wundrack, E.-D. Gilles, S. Klamt, T. Kähne and M. Naumann (2008). “Host-pathogen systems biology: logical modelling of hepatocyte growth factor and Helicobacter pylori induced c-Met signal transduction.” BMC Systems Biology 2(1): 4.

Franke, R., M. Müller, N. Wundrack, E. D. Gilles, S. Klamt, T. Kähne and M. Naumann (2008). “Host-pathogen systems biology: logical modelling of hepatocyte growth factor and Helicobacter pylori induced c-Met signal transduction.” BMC Syst Biol 2: 4.

Furge, K. A., Y. W. Zhang and G. F. Vande Woude (2000). “Met receptor tyrosine kinase: enhanced signaling through adapter proteins.” Oncogene 19(49): 5582–5589.

Hayashida, A., S. Amano and P. W. Park (2011). “Syndecan-1 promotes Staphylococcus aureus corneal infection by counteracting neutrophil-mediated host defense.” J Biol Chem 286(5): 3288–3297.

Horridge, D. N., A. A. Begley, J. Kim, N. Aravindan, K. Fan and M. H. Forsyth (2017). “Outer inflammatory protein a (OipA) of Helicobacter pylori is regulated by host cell contact and mediates CagA translocation and interleukin-8 response only in the presence of a functional cag pathogenicity island type IV secretion system.” Pathog Dis 75(8).

Imamura, R. and K. Matsumoto (2017). “Hepatocyte growth factor in physiology and infectious diseases.” Cytokine 98: 97–106.

Ito, N., H. Tsujimoto, H. Ueno, Q. Xie and N. Shinomiya (2020). “Helicobacter pylori-Mediated Immunity and Signaling Transduction in Gastric Cancer.” Journal of Clinical Medicine 9(11):

Ito, N., H. Tsujimoto, H. Ueno, Q. Xie and N. Shinomiya (2020). “Helicobacter pylori-Mediated Immunity and Signaling Transduction in Gastric Cancer.” J Clin Med 9(11).

Jumper, J., R. Evans, A. Pritzel, T. Green, M. Figurnov, O. Ronneberger, K. Tunyasuvunakool, R. Bates, A. Zidek, A. Potapenko, A. Bridgland, C. Meyer, S. A. A. Kohl, A. J. Ballard, A. Cowie, B. Romera-Paredes, S. Nikolov, R. Jain, J. Adler, T. Back, S. Petersen, D. Reiman, E. Clancy, M. Zielinski, M. Steinegger, M. Pacholska, T. Berghammer, S. Bodenstein, D. Silver, O. Vinyals, A. W. Senior, K. Kavukcuoglu, P. Kohli and D. Hassabis (2021). “Highly accurate protein structure prediction with AlphaFold.” Nature 596(7873): 583–589.

Kalali, B., R. Mejías-Luque, A. Javaheri and M. Gerhard (2014). “H. pylori virulence factors: influence on immune system and pathology.” Mediators Inflamm 2014: 426309.

Kao, C. Y., B. S. Sheu and J. J. Wu (2016). “Helicobacter pylori infection: An overview of bacterial virulence factors and pathogenesis.” Biomed J 39(1): 14–23.

Kurien, B. T. and R. H. Scofield (2012). “Extraction of proteins from gels: a brief review.” Methods Mol Biol 869: 403–405.

Lima, V. P., M. A. de Lima, M. V. Ferreira, M. A. Barros and S. H. Rabenhorst (2010). “The relationship between Helicobacter pylori genes cagE and virB11 and gastric cancer.” Int J Infect Dis 14(7): e613–617.

Liu, C. L. and G. P. Shi (2019). “Calcium-activated chloride channel regulator 1 (CLCA1): More than a regulator of chloride transport and mucus production.” World Allergy Organ J 12(11): 100077.

Loewen, M. E. and G. W. Forsyth (2005). “Structure and function of CLCA proteins.” Physiol Rev 85(3): 1061–1092.

Mariani, V., M. Biasini, A. Barbato and T. Schwede (2013). “lDDT: a local superposition-free score for comparing protein structures and models using distance difference tests.” Bioinformatics 29(21): 2722–2728.

Martin, E. R., A. Barbieri, R. C. Ford and R. C. Robinson (2020). “In vivo crystals reveal critical features of the interaction between cystic fibrosis transmembrane conductance regulator (CFTR) and the PDZ2 domain of Na(+)/H(+) exchange cofactor NHERF1.” J Biol Chem 295(14): 4464–4476.

Matsuo, Y., Y. Kido and Y. Yamaoka (2017). “Helicobacter pylori Outer Membrane Protein-Related Pathogenesis.” Toxins (Basel) 9(3).

Meyer-ter-Vehn, T., A. Covacci, M. Kist and H. L. Pahl (2000). “Helicobacter pylori Activates Mitogen-activated Protein Kinase Cascades and Induces Expression of the Proto-oncogenes c-fos and c-jun *.” Journal of Biological Chemistry 275(21): 16064–16072.

Min, L., Y. Zhao, S. Zhu, X. Qiu, R. Cheng, J. Xing, L. Shao, S. Guo and S. Zhang (2017). “Integrated Analysis Identifies Molecular Signatures and Specific Prognostic Factors for Different Gastric Cancer Subtypes.” Translational Oncology 10(1): 99–107.

Mirdita, M., K. Schutze, Y. Moriwaki, L. Heo, S. Ovchinnikov and M. Steinegger (2022). “ColabFold: making protein folding accessible to all.” Nat Methods 19(6): 679–682.

Niemann, H. H., V. Jäger, P. J. Butler, J. van den Heuvel, S. Schmidt, D. Ferraris, E. Gherardi and D. W. Heinz (2007). “Structure of the human receptor tyrosine kinase met in complex with the Listeria invasion protein InlB.” Cell 130(2): 235–246.

Oleastro, M. and A. Ménard (2013). “The Role of Helicobacter pylori Outer Membrane Proteins in Adherence and Pathogenesis.” Biology (Basel) 2(3): 1110–1134.

Pantzar, M., A. Ljungh and T. Wadström (1998). “Plasminogen binding and activation at the surface of Helicobacter pylori CCUG 17874.” Infect Immun 66(10): 4976–4980.

Peek, R. M., Jr. and M. J. Blaser (2002). “Helicobacter pylori and gastrointestinal tract adenocarcinomas.” Nat Rev Cancer 2(1): 28–37.

Polat, A. N., Ö. Karayel, S. H. Giese, B. Harmanda, E. Sanal, C. K. Hu, B. Y. Renard and N. Özlü (2015). “Phosphoproteomic Analysis of Aurora Kinase Inhibition in Monopolar Cytokinesis.” J Proteome Res 14(9): 4087–4098.

Posselt, G., S. Backert and S. Wessler (2013). “The functional interplay of Helicobacter pylori factors with gastric epithelial cells induces a multi-step process in pathogenesis.” Cell Commun Signal 11: 77.

Qiao, S., Q. Luo, Y. Zhao, X. C. Zhang and Y. Huang (2014). “Structural basis for lipopolysaccharide insertion in the bacterial outer membrane.” Nature 511(7507): 108–111.

Ritter, B., J. Philie, M. Girard, E. C. Tung, F. Blondeau and P. S. McPherson (2003). “Identification of a family of endocytic proteins that define a new alpha-adaptin ear-binding motif.” EMBO Rep 4(11): 1089–1095.

Svensson, H., M. Hansson, J. Kilhamn, S. Backert and M. Quiding-Järbrink (2009). “Selective Upregulation of Endothelial E-Selectin in Response to <i>Helicobacter pylori</i>-Induced Gastritis.” Infection and Immunity 77(7): 3109–3116.

Szulc-Dąbrowska, L., M. Bossowska-Nowicka, J. Struzik and F. N. Toka (2020). “Cathepsins in Bacteria-Macrophage Interaction: Defenders or Victims of Circumstance?” Frontiers in Cellular and Infection Microbiology 10.

Tunyasuvunakool, K., J. Adler, Z. Wu, T. Green, M. Zielinski, A. Žídek, A. Bridgland, A. Cowie, C. Meyer, A. Laydon, S. Velankar, G. J. Kleywegt, A. Bateman, R. Evans, A. Pritzel, M. Figurnov, O. Ronneberger, R. Bates, S. A. A. Kohl, A. Potapenko, A. J. Ballard, B. Romera-Paredes, S. Nikolov, R. Jain, E. Clancy, D. Reiman, S. Petersen, A. W. Senior, K. Kavukcuoglu, E. Birney, P. Kohli, J. Jumper and D. Hassabis (2021). “Highly accurate protein structure prediction for the human proteome.” Nature 596(7873): 590–596.

van Zundert, G. C. P., J. Rodrigues, M. Trellet, C. Schmitz, P. L. Kastritis, E. Karaca, A. S. J. Melquiond, M. van Dijk, S. J. de Vries and A. Bonvin (2016). “The HADDOCK2.2 Web Server: User-Friendly Integrative Modeling of Biomolecular Complexes.” J Mol Biol 428(4): 720–725.

Wu, J. H., B. J. Billings and D. F. Balkovetz (2001). “Hepatocyte Growth Factor Alters Renal Epithelial Cell Susceptibility to Uropathogenic Escherichia coli.” Journal of the American Society of Nephrology 12(12): 2543–2553.

Xie, C., Z. Yang, Y. Hu, X. Cao, J. Chen, Y. Zhu and N. Lu (2017). “Expression of c-Met and hepatocyte growth factor in various gastric pathologies and its association with Helicobacter pylori infection.” Oncol Lett 14(5): 6151–6155.

Xu, C., D. M. Soyfoo, Y. Wu and S. Xu (2020). “Virulence of Helicobacter pylori outer membrane proteins: an updated review.” Eur J Clin Microbiol Infect Dis 39(10): 1821–1830.

Zhang, Y., X. G. Fan, R. Chen, Z. Q. Xiao, X. P. Feng, X. F. Tian and Z. H. Chen (2005). “Comparative proteome analysis of untreated and Helicobacter pylori-treated HepG2.” World J Gastroenterol 11(22): 3485–3489.

Zhu, W., A. Shenoy, P. Kundrotas and A. Elofsson (2023). “Evaluation of AlphaFold-Multimer prediction on multi-chain protein complexes.” Bioinformatics 39(7).

